# Intimate functional interactions between TGS1 and the Smn complex revealed by an analysis of the Drosophila eye development

**DOI:** 10.1101/2020.02.06.936724

**Authors:** Paolo Maccallini, Francesca Bavasso, Livia Scatolini, Elisabetta Bucciarelli, Gemma Noviello, Veronica Lisi, Valeria Palumbo, Simone D’Angeli, Stefano Cacchione, Giovanni Cenci, Laura Ciapponi, James G. Wakefield, Maurizio Gatti, Grazia Daniela Raffa

**Author notes:** Equal contribution (Paolo Maccallini, Francesca Bavasso). Correspondence: Grazia Daniela Raffa, Maurizio Gatti.

## Abstract

Trimethylguanosine synthase 1 (TGS1) is a conserved enzyme that mediates formation of the trimethylguanosine cap on several RNAs, including snRNAs and telomerase RNA. Previous studies have shown that TGS1 binds the Survival Motor Neuron (SMN) protein, whose deficiency causes spinal muscular atrophy (SMA). In addition, TGS1 depletion results in increased hTR levels and telomere elongation in human cells. Here, we analyzed the roles of the *Drosophila* orthologs of the human *TGS1* and *SMN* genes. We show that the *Drosophila* TGS1 protein (dTgs1) physically interacts with all subunits of the *Drosophila* Smn complex (Smn, Gem2, Gem3, Gem4 and Gem5), and that a human *TGS1* transgene rescues the mutant phenotype caused by *dTgs1* loss. We demonstrate that both *dTgs1* and *Smn* are required for viability of retinal progenitor cells and that downregulation of these genes leads to a reduced eye size. Importantly, overexpression of *dTgs1* partially rescues the eye defects caused by Smn depletion, and vice versa. These results suggest that the *Drosophila* eye model can be exploited for screens aimed at the identification of genes and drugs that modify the phenotypes elicited by Tgs1 and Smn deficiency. These modifiers could help to devise new therapies for SMA and diseases caused by telomerase insufficiency.

## Introduction

Trimethylguanosine synthase 1 (TGS1) catalyzes conversion of the 5’ mono-methylguanosine cap (MMG) of RNA into a trimethylguanosine cap (TMG). TGS1 is evolutionarily conserved and mediates hypermethylation of a variety of Pol II-dependent RNAs, including small nuclear (sn) RNAs, small nucleolar (sno) RNAs, telomerase RNA and selenoprotein mRNAs [1-4]. TGS1 is not essential for viability in *Saccharomyces cerevisiae, Schizosaccharomyces pombe* or *Arabidopsis thaliana*, but loss of TGS1 renders both *S. cerevisiae* and *A. thaliana* sensitive to cold [2,5,6]. In contrast, loss of TGS1 causes larval lethality in *Drosophila melanogaster* [7-9], and leads to early embryonic lethality in mice [10], indicating that cap hypermethylation has an essential role in animal development.

Studies in human cells have defined the role of TGS1 in maturation and trafficking of small RNAs. In human cells, there are two TGS1 isoforms, a long isoform (TGS1-LF) that contains the methyltransferase domain of the enzyme, and a short (TGS1-SF) isoform that consists only of the C terminus of the protein. TGS1-LF is present in both the cytoplasm and the nuclear Cajal bodies (CBs) and regulates trafficking of both snoRNAs and snRNAs; TGS1-SF is restricted to the CBs where it specifically interacts with snoRNAs [1]. In the nucleus, the monomethylated 5’ cap of snRNAs binds the cap-binding complex (CBC) that mediates their export to the cytoplasm through an interaction with the CRM1 and PHAX export factors [11]. Once in the cytoplasm, snRNAs associate with the Sm protein complex that physically binds TGS1 through its SmB component [2,12]. The seven Sm core proteins assemble into a heteroheptameric donut-shaped multiprotein structure that binds the U1, U2, U4 and U5 snRNAs, forming four of the five snRNP subunits of the spliceosome [13]. The assembly of the Sm-snRNA particles is chaperoned by the survival of motor neurons (SMN) complex, which includes SMN, Gemin2-8 and Unrip/STRAP [14]. Following the snRNA interaction with the Sm and SMN complexes, TGS1 hypermethylates the MMG cap of snRNAs, and the TMG-snRNPs are reimported into the nucleus [1,2].

TGS1 has also been implicated in the regulation of the telomerase RNA moiety. In *S. cerevisiae*, TGS1catalyzes TMG cap formation on TLC1 (the RNA component of *S. cerevisiae* telomerase), and its loss causes telomere lengthening and an increase in telomere silencing [15]. TGS1 also mediates TER1 (the RNA of *S. pombe* telomerase) hypermethylation *in S. pombe*. However, loss of TGS1 in this yeast affects TER1 processing and stability, resulting in telomere shortening [16]. We have recently found that TGS1 hypermethylates human telomerase RNA (hTR) and that TGS1 deficiency increases both the mature hTR level and telomerase activity, leading to telomere elongation [4]. Thus, while TGS1 mediates the formation of a TMG cap in both yeasts ad humans, the effects of this post-transcriptional modification on telomerase regulation appear to be species-specific.

In humans, an impairment of *SMN* function causes Spinal Muscular Atrophy (SMA), a devastating recessive disorder characterized by motor neuron loss, progressive paralysis and death [17]. The human genome harbors two *SMN* genes, but *SMN2* does not produce a sufficient amount of protein to compensate for homozygous *SMN1* mutations found in SMA patients. Although the exact mechanisms through which SMN deficiency disrupts motor neuron function have not been fully elucidated, one of the most accredited hypotheses is that loss of the SMN reduces the snRNP levels in neural cells, resulting in splicing defects in mRNAs with critical roles in motor neuron function and maintenance [18-22]. SMN targets include mRNAs encoding two negative regulators of the abundance of p53 [23], a key driver of motor neuron death [24]. Another SMN target, *Stasimon*, plays a dual role by preserving the function of the sensory-motor circuit and by restricting phosphorylation-mediated p53 activation [25]. SMN also plays splicing-independent functions that are thought to contribute to the SMA phenotype. These functions include regulation of axonal localization and transport of mRNPs [26-29], proper development and regeneration of the neuromuscular junction [30], neural stem cell division and differentiation [31,32], integrity and function of skeletal muscle [33], and prevention of transcriptional stress and DNA damage [34-36].

In *Drosophila* there is only one *TGS1* gene that is part of a bicistronic locus that also includes *modigliani* (*moi*), a gene that encodes a telomeric protein required to prevent telomere fusion [8,9]. *Drosophila TGS1* (henceforth *dTgs1*) is an essential gene [8,9] that mediates snRNA hypermethylation [7] and interacts with Gemin3 subunit of the SMN complex in the two-hybrid assay [37]. It has been reported that dTgs1 depletion affects fly motor behavior, eliciting phenotypes reminiscent of those observed in Smn-deficient animals [8,37]. Here, we show that dTgs1 physically interacts with all subunits of the *Drosophila* Smn complex, and that a human *TGS1* transgene fully rescues the lethal phenotype of d*Tgs1* null mutants. In addition, we demonstrate that both *dTgs1* and *Smn* are required for *Drosophila* eye development, and that the two genes cooperate to ensure viability of retinal progenitor cells. Importantly, we show that overexpression of *dTgs1* partially rescues the eye defects caused by *Smn* deficiency, while *Smn* overexpression ameliorates the eye phenotype elicited by dTgs1 depletion. Thus our work establishes a new *Drosophila* model that can be exploited for a variety of functional studies on both *dTgs1* and *Smn*.

## Materials and Methods

### Drosophila strains and transgenic constructs

The *moi*^*2*^, *dTgs1*^*R1*^ and *dTgs1*^*R2*^ mutant alleles were generated by CRISPR/Cas9 genome editing. The *moiCRISPR guide (*CTCTCGAGGTAGAGGCTTC) was cloned into pCFD3-dU6:3gRNA (Addgene plasmid # 49410; http://n2t.net/addgene:49410; [38]. The *Tgs1CRISPR* guide (*TATCGAGGTGGTTTCGTCGG)* was cloned into pU6-BbsI-chiRNA (Addgene plasmid # 45946; http://n2t.net/addgene:45946; [39]. The *dTgs1* ^*m1*^ mutant strain is the *moi*^*1*^ mutant strain [9]. The *UAS-GFP-dTgs1* strain carries the pPGW*-Tgs1* construct, generated by cloning the *dTgs1* CDS into the pPGW destination vector (Stock number 1077, Drosophila Genomics Resource Center, supported by NIH grant 2P40OD010949), using the Gateway technology (Thermo Fisher Scientific); The *UAS-hTGS1* and *UAS-hTGS1*^*CD*^ transgenes carry full-length human *TGS1* genes cloned into the pUAST-attB vector [40]; the *hTGS1*^*CD*^ gene was generated by site-directed mutagenesis using the Gibson Assembly® Master Mix to produce a cDNA encoding the amino acid substitutions indicated in figure 5A. Transgenic flies were obtained by injecting the constructs into *y*^*1*^ *v*^*1*^; *P{CaryP}attP40* (2L, 25C6). The *tub-Smn*-GFP strain used for AP/MS experiments carries a construct the eGFP sequences fused in-frame with the 3′ end of the *Smn* CDS under control of the tubulin promoter, cloned into the pJZ4 vector [41,42]. All embryo injections were carried out by BestGene (Chino Hills, CA, USA).

The *tub-Smn-FLAG* bearing strain was described in [42].The *moi*^*+*^, *moi-dTgs1*^*FL*^ and *moi-dTgs1*^*CD*^ carrying stocks were described in [9]. The UAS-d*Tgs1RNAi* stock P{GD14932}v29503, was obtained from the Vienna Drosophila Resource Center (VDRC, www.vdrc.at) [43]. The *UAS-SmnRNAi* stock P{TRiP.HMC03832}attP40, along with the P{w^+mC^ = Act5C-Gal4}25FO1, the P{GawB}how24B, the P{nSyb-Gal4.S}3, the P{GawB}elavC155, and P{Gal4-ey.H}3-8 drivers were obtained from the Bloomington Drosophila Stock Center (NIH P40OD018537). *The Smn*^*X7*^ mutant allele, a deficiency that uncovers most of the *Smn* coding sequence, and is gift from Dr. Artavanis-Tsakonas [44]. The Oregon-R strain was used as a wild type control. All flies were reared according to standard procedures at 25 °C. Lethal mutations were balanced over either *TM6B, Hu, Tb* or *CyO-TbA, Cy, Tb* [45]. *All* genetic markers and special chromosomes are described in detail in FlyBase (http://www.flybase.org).

### GFP-TRAP-A isolation of GFP-Tgs1 and Smn-GFP

For AP/MS experiments we used embryos from females expressing *GFP-dTgs1* from the pPGW*-Tgs1* construct induced by *Act-Gal4*, or *Smn*-GFP from a tub-*Smn*-GFP transgene (see above for description of the transgenic stocks). Batches of 0-3 h old embryos laid by cages of 1-10 days-old flies were dechorionated, weighed, flash frozen in liquid nitrogen and stored at -80°C. For MS analysis, the following procedure was undertaken: ∼0.4 g of frozen embryos were homogenized in 1.5 ml of C buffer (50 mM HEPES [pH 7.4], 50 mM KCl, 1 mM MgCl2, 1 mM EGTA, 0.1% IGEPAL CA-630, protease inhibitors (Roche). Extract was clarified through centrifugation at 10,000 g for 10 min, 100,000 g for 30 min, and 100,000 g for a further 10 min. Clarified extract was incubated with 15 μl GFP-TRAP-A beads or blocked agarose beads (bab-20) (Chromotek) equilibrated in C Buffer for 2 h at 4°C. Tgs1 GFP/GFP-TRAP-A beads were then washed 4 times with ice-cold C buffer and stored at -20°C.

### Mass spectrometric analysis

Mass spectrometric analysis was undertaken by the Bristol Proteomics Facility (*http://www.bristol.ac.uk/biomedical-sciences/research/facilities/proteomics/*), Samples were run ∼1 cm into the separating region of an SDS-PA gel, cut as a single slice and subjected to in-gel tryptic digestion using a DigestPro automated digestion unit (Intavis Ltd.). The resulting peptides were fractionated using a Dionex Ultimate 3000 nanoHPLC system in line with an LTQ-Orbitrap Velos mass spectrometer (Thermo Scientific). In brief, peptides in 1% (vol/vol) formic acid were injected onto an Acclaim PepMap C18 nano trap column (Dionex). After washing with 0.5% (vol/vol) acetonitrile 0.1% (vol/vol) formic acid peptides were resolved on a 250 mm × 75 μm Acclaim PepMap C18 reverse phase analytical column (Dionex) over a 150 min organic gradient, using 7 gradient segments (1-6% solvent B over 1 min, 6 15% B over 58 min, 15-32% B over 58 min, 32-40% B over 3 min, 40-90% B over 1 min, held at 90% B for 6 min and then reduced to 1% B over 1 min) with a flow rate of 300 nl min-1. Solvent A was 0.1% formic acid and Solvent B was aqueous 80% acetonitrile in 0.1% formic acid. Peptides were ionized by nano-electrospray ionization at 2.1 kV using a stainless steel emitter with an internal diameter of 30 μm (Thermo Scientific) and a capillary temperature of 250°C. Tandem mass spectra were acquired using an LTQ-Orbitrap Velos mass spectrometer controlled by Xcalibur 2.1 76 software (Thermo Scientific) and operated in data-dependent acquisition mode. The Orbitrap was set to analyse the survey scans at 60,000 resolution (at m/z 400) in the mass range m/z 300 to 2000 and the top twenty multiply charged ions in each duty cycle selected for MS/MS in the LTQ linear ion trap. Charge state filtering, where unassigned precursor ions were not selected for fragmentation, and dynamic exclusion (repeat count, 1; repeat duration, 30 s; exclusion list size, 500) was used. Fragmentation conditions in the LTQ were as follows: normalized collision energy, 40%; activation q, 0.25; activation time, 10 ms; and minimum ion selection intensity, 500 counts. The raw data files were processed and quantified using Proteome Discoverer software v1.2 (Thermo Scientific) and searched against the dmel-all translation-r5.47 database using the SEQUEST (Ver. 28 Rev. 13) algorithm. Peptide precursor mass tolerance was set at 10ppm, and MS/MS tolerance was set at 0.8Da. Search criteria included carbamidomethylation of cysteine (+57.0214) as a fixed modification and oxidation of methionine (+15.9949) as a variable modification. Searches were performed with full tryptic digestion and a maximum of 1 missed cleavage was allowed. The reverse database search option was enabled and all peptide data was filtered to satisfy false discovery rate (FDR) of 5%.

### Bioinformatics filtering of MS data

For stringent filtering, MS results were filtered by removing protein IDs with (i) single peptide hits, (ii) <20% peptide:protein coverage and (iii) overall MS Scores of <50. Remaining IDs were cross-referenced against an accumulated database of “false-positives”; MS data used as negative controls were accumulated from eight independent control GFP-TRAP-A experiments, each using extracts from ∼0.4 g 0-3 h embryos expressing GFP fusions to proteins in which a bait protein was not precipitated, as described by [46]. Any protein ID that was not identified in negative control list and with a score above 50 are presented in Figure 2. The full datasets are provided in Tables S1 and S2.

### Larval Locomotion Analysis

Larval locomotion analysis was described previously [42]. Briefly, locomotor activity was measured by counting the number of peristaltic contractions per minute of third instar larvae on a surface of a 1% agarose gel in a Petri dish; measurements were repeated ten times. To obtain unbiased measurement of locomotion parameters, larvae were blind-tested by three experimenters. The significance of multiple comparisons was evaluated with One Way Analysis of Variance. The Tukey’s test was performed as Post-Hoc Test to determine the significance between every single group. (P<0.01 was considered significant).

### Microscopic analysis of Eye-antennal imaginal discs

Third instar larval eye-antennal imaginal discs were dissected in ice cold PBS and fixed in 4% paraformaldehyde in PBS at room temperature for 30 min. After fixation, the tissues were washed in PBS 0,3% Triton X-100 (PBS-T) for 3 x 20 min and blocked with PBS-T and 5% BSA for 30 min. Discs were incubated overnight at 4°C with anti-Cleaved Caspase3 (1:300; Cell Signaling Technology) and mouse anti-Elav [1:100; Developmental Studies Hybridoma Bank (DSHB)], washed 3 x 20 min in PBS-T, and then incubated for 1 hr at room temperature with Cy3-conjugated anti-rabbit (1:300, Life Technologies) and FITC-conjugated anti-mouse (1:100, Jackson Laboratories). All samples were mounted in Vectashield with DAPI (Vector) to stain DNA and reduce fluorescence fading. Images were acquired using an Axio Imager M2 fluorescence microscope (Zeiss, Germany). Image z-stacks were acquired with an Axiocam 512 (Zeiss) monochromatic camera and Apotome 2 (Zeiss); for each image15 z-stack were acquired at 0,5 micrometer Z step. All images were acquired with the same parameters. Fluorescence signals were quantified using Zen 2.5 Pro software (Zeiss, Germany). Measurements of the eye-antennal disc areas and the ELAV stained areas were performed with the Zen 2.5 Pro software (Zeiss, Germany) on Maximum intensity projections z-stacks. Caspase3 positive foci were also quantified using Zen 2.5 Pro software. Fluorescence threshold was setup starting from the basal fluorescence, and spots were considered positive starting from 3 times the basal fluorescence. We considered only spots with areas greater than 1 μm but less than 60 μm and calculated the mean value their fluorescence signal. Statistical significance was calculated using the Kruskal-Wallis test with GraphPad Prism.

### Evaluation of the eye phenotype

The eye size of RNAi flies was evaluated by comparison with the wild type eye (Oregon-R). Abnormal eyes were assigned to one of the following classes: class 100 that comprises eyes of normal size, and classes 25, 50, and 75 that include eyes with sizes < 25%, < 50%, and < 75% the size of the wild type eye, respectively. Eyes were classified by visual inspection performed independently by at least two researchers. When classification was not clear-cut, the eye was assigned to the higher class in the evaluation of RNAi phenotype without rescue construct (e.g., an eye of dubious class 50 was assigned to class 75), and to the lower class in the presence of a rescue construct.

### Generation of an anti-dTgs1 antibody

To generate an anti-dTgs1 rabbit polyclonal antiserum, the dTgs1 sequence coding for aa 335-466 was cloned into the pGEX vector to produce a peptide fused to GST. The GST-fusion peptide was expressed in BL21-CodonPlus Competent Cells-Agilent, and purified by incubating crude lysates with glutathione sepharose 4B (Amersham), as recommended by the manufacturer. Rabbit Immunization was carried out at Agro-Bio Services (www.agro-bio.com). The specificity of the antiserum was confirmed by Western blotting on protein extracts from homozygous *dTgs1* null mutants and control flies (Figure 3C). In control flies, the antiserum detects a protein of the expected MW for dTgs1 (∼ 60 KDa) that is strongly reduced in mutants.

### Western Blotting

Protein extracts from 15 third instar larval brains, lysed in sample buffer were fractionated by SDS-PAGE and transferred to nitrocellulose membrane. Membranes were probed with rabbit anti-Tgs1 antiserum (1:5000; this study), mouse anti-tubulin (1:20000; Sigma-Aldrich), anti-beta Actin (1:100000, Abcam 49900 [AC-15], HRP), rabbit anti-human TGS1 (1:1500, Bethyl Laboratories Cat#A300-814A, lot 1), rabbit anti-GFP antibody (1:1000; Torrey Pines Biolabs, TP401). These primary antibodies were detected with HRP-conjugated anti-mouse or anti-rabbit (1:5000; GE HealthCare), using the SuperSignal™ West Pico chemiluminescent substrate (Thermo Fisher Scientific); images were acquired with Chemidoc (Biorad) and quantified using the QuantityOne image analysis software (Biorad).

## Results

### Analysis of the *Drosophila moi-dTgs1* bicistronic locus

*dTgs1* is part of a bicistronic locus that also includes *moi*, which encodes a protein required for telomere capping (Figure 1A). Previous work based on the analysis of rescue experiments of mutations and deficiencies involving the *moi*-*dTgs1* bicistronic locus has suggested that the two genes play independent functions [8,9]. To provide a definitive proof for this independency we generated several *moi* and *dTgs1* mutations using CRISPR/Cas9 mutagenesis. The *moi* mutation we characterized (*moi*^*2*^) contains a frameshift mutation resulting in an early stop codon (Figure 1A, B). Homozygous *moi*^*2*^ animals are lethal, dying in third instar larvae/early pupal stages; they exhibit ∼ 3 telomeric fusions per larval brain cell (n = 200) and are fully rescued by a *moi+* transgene constitutively driven by the tubulin promoter (Figure 1B). The two CRISPR/Cas9-induced *Tgs1* mutations (*dTgs1*^*R1*^ and *dTgs1*^*R2*^) result in early stop codons in the *dTgs1* sequence (Figure 1B). To characterize these mutations we generated a polyclonal antibody against a C-terminal fragment of dTgs1 (See Materials and Methods). Western blotting analysis showed that *dTgs1*^*R1*^ and *dTgs1*^*R2*^ homozygotes exhibit a strongly reduced dTgs1 band compared to wild type (Figure 1C). Homozygotes for each of these mutations die as second instar larvae but are viable in the presence of either a ubiquitously expressed *dTgs1*^*FL*^ transgene including both *moi*^*+*^ and *dTgs1*^*+*^ or an inducible *UAS-GFP-dTgs1* transgene driven by *actin-GAL4* (Figure 1B). In contrast, *dTgs1*^*R1*^ and *dTgs1*^*R2*^ homozygotes, constitutively expressing the wild type *moi* transgene (*moi*^*+*^), still die as second instar larvae (Figure 1B).

**Figure 1.**
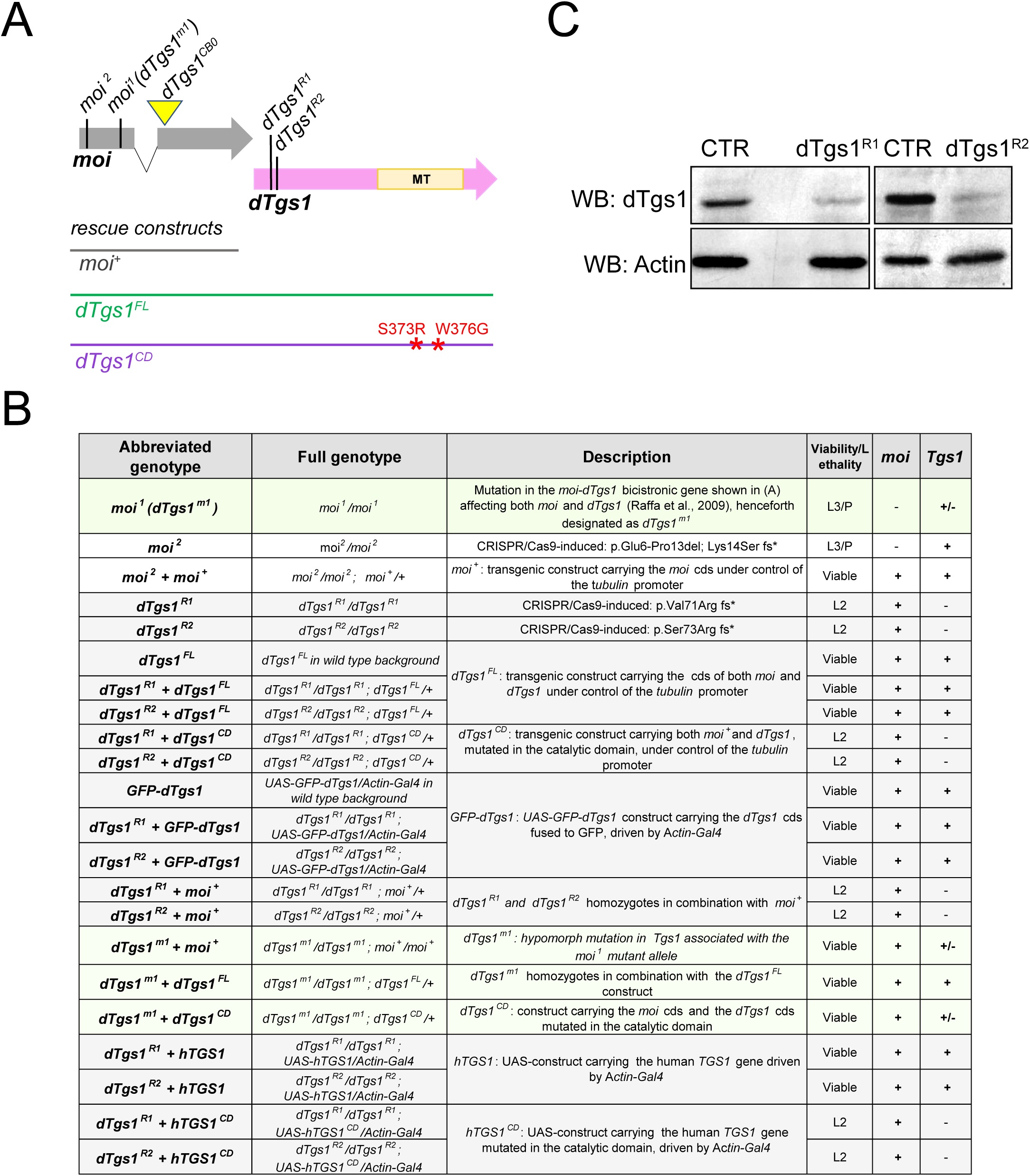
*dTgs1* mutations and mutant combinations used in this study. (A) Schematic representation of the *Drosophila* bicistronic locus *CG31241*, which produces a unique transcript containing two ORFs. The first ORF (grey) is interrupted by a small intron and encodes the telomere capping protein Moi; the second ORF (pink) encodes *Drosophila Tgs1 (dTgs1)*. The vertical lines and the yellow triangle indicate the positions of the point and insertional (triangle) mutations described in B. The horizontal lines represent the genomic fragments fused with the tubulin promoter used as rescue constructs. FL: full length; CD: catalytic dead; MT: methyl-transferase domain. (B) Transgenic constructs and mutant combinations used in this study. For each combination is reported the viability or the lethal stage (L2, lethal in second larval instar; L3, lethal in third larval instar; P lethal in the pupal stage), and the *moi* and *dTgs1* proficiency (+, normal; +/- reduced; - virtually null). (C) Western blots of extracts from second instar larvae homozygous for either the *dTgs1*^*R1*^ or the *dTgs1*^*R2*^ mutation, showing a strong reduction in dTgs1 compared to wild type larvae at the same developmental stage (CTR). The dTgs1 protein in mutant larvae is likely to be maternally supplied. Actin is a loading control.

Importantly, the *moi*^*2*^*/dTgs1*^*R1*^ and *moi*^*2*^/*dTgs1*^*R1*^ heterozygous flies were fully viable, indicating that the protein products of the *moi* and *dTgs1* genes are functionally independent. Collectively, these genetic and cytological analyses indicate that *dTgs1* is an essential *Drosophila* gene that functions independently of *moi*. Previous work indicated that overexpression of a *UAS-Tgs1*-3XHA transgene, driven by the ubiquitous *daughterless-Gal4* driver, is lethal [37]. In contrast, we found that overexpression of *UAS-GFP-dTgs1* driven by the *Actin-Gal4* ubiquitous driver is fully viable (Figure 1B). Also viable are flies carrying either one or two copies of the *dTgs1*^*FL*^ construct (*tubulin-moi-dTgs1*) (Figure 1B). We do not understand the reason for this discrepancy, which might depend on the type of UAS construct used, the driver, or both.

### dTgs1 physically interacts with the Smn complex

Previous work in human cells has shown that snRNAs bind the Sm proteins in the cytoplasm, with the assistance of the SMN complex that acts as a molecular chaperone [47]. Following the interaction with the Sm and SMN complexes, the snRNAs are hypermethylated by TGS1 and reimported into the nucleus [1,2,48]. During this process TGS1 directly interacts with the SMN protein and the SmB component of the Sm complex [12]. These findings prompted us to investigate whether dTgs1 interacts with the fly Smn complex. To identify the dTgs1 interacting partners we incubated extracts from 0-3 hour old embryos expressing GFP-dTgs1 with GFP-TRAP-A, and subjected them to affinity purification-mass spectrometry (AP-MS). The fly line used for this experiment carried a *UAS-GFP-dTgs1* transgene on one of its second chromosomes and an *actin-GAL4* driver on the other, and was homozygous for the null mutation d*Tgs1*^*R2*^ such that the flies of this line express GFP-tagged dTgs1 but not the endogenous dTgs1 protein. GFP-dTgs1 was efficiently purified (Figure 2A) and was the most abundant protein in precipitates (Figure 2B and Table S1). To select bona fide dTgs1-binding partners we applied stringent filtering against a database of non-specific interactors as described by [46]; we selected only proteins IDs that were not found in the negative control and with a score above 50, and ranked them by mean area. Using this criterion, we identified 11 specific dTgs1 interactors (Figure 2B) that include Smn and four Gemins: Gem2, Gem3 and the protein products of the duplicated genes *Gem4a* (*Glos* or *CG2941*) and *Gem4b* (*CG32786*) [49,50]. Peptides corresponding to Gem5 were also detected in precipitates, but with an overall score well below our cut-off (Figure 2B; Table S1). Thus, dTgs1 co-purifies with most components of the *Drosophila* Smn complex (Figure 2B; Table S1)

To confirm and extend these results we also performed AP-MS from 0-3 hours embryos expressing Smn-GFP. For this experiment we used embryos from mothers carrying two wild type copies of the *Smn* gene and homozygous for an *Smn-GFP* transgene placed under the control of the tubulin promoter [42]. Using the same criterion described above for dTgs1, Smn was the most abundant protein in precipitates, along with 11 specific interactors, again including Gem2, Gem3, Gem4a and Gem4b (Figure 2C, Table S2). In addition, Smn-GFP co-purified Lsm11, the Cap binding protein (Cbp80) and dTgs1 (Figure 2C). Gem5 and Lsm10 were also present in precipitates but fell just outside of our stringent cut-off criteria (Figure 2C).These findings are in good agreement with recent AP-MS results obtained from *Drosophila* embryos expressing Smn-FLAG [49]. Collectively, our results provide strong evidence that dTgs1 physically associates with the *Drosophila* Smn complex in vivo, suggesting a parallel functional interaction.

**Figure 2.**
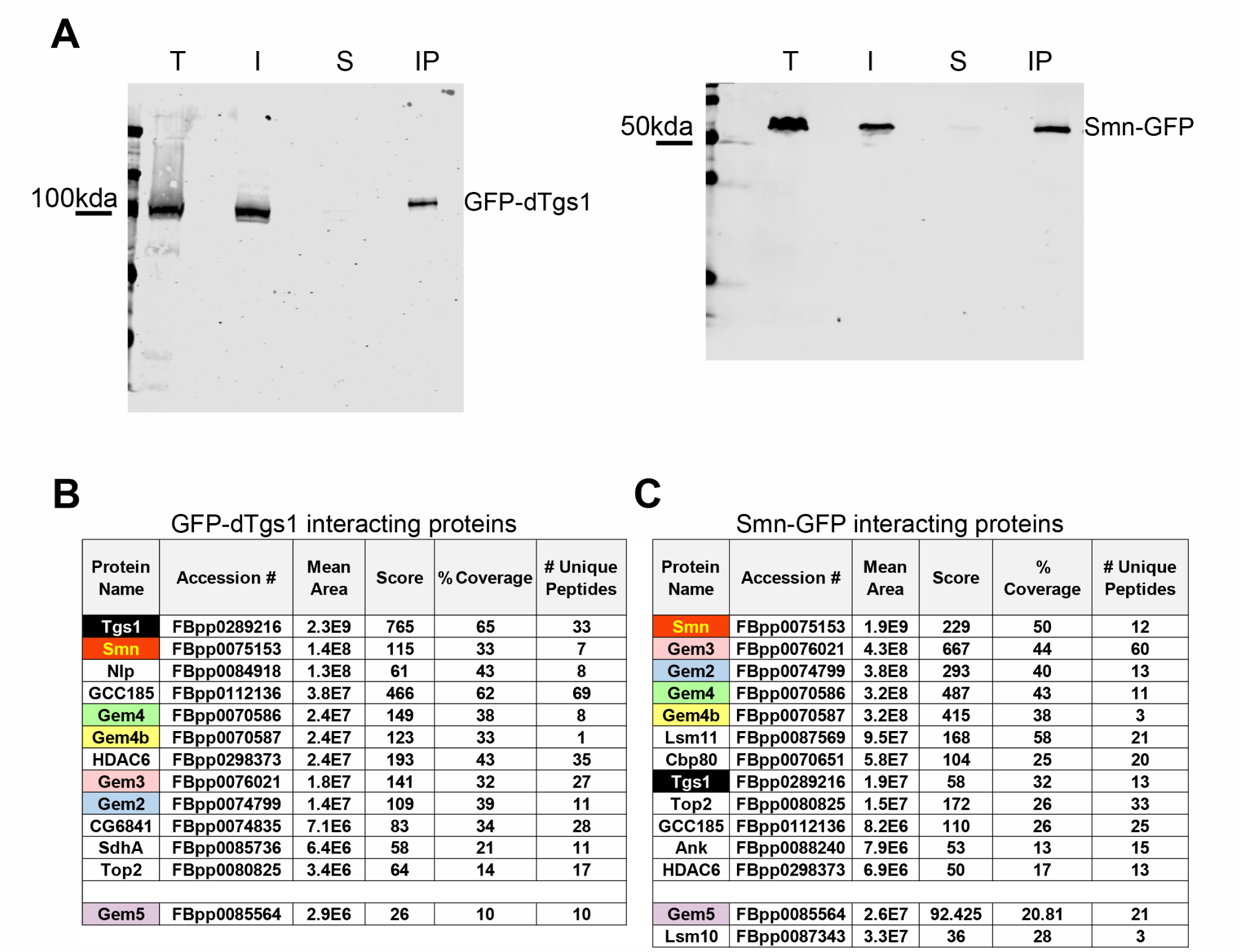
Physical interactions between dTgs1 and the SMN complex. Proteins co-purifying with dTgs1 or Smn were identified by affinity purification (AP) using GFP-TRAP beads, followed by mass spectrometry (MS) and stringent filtering (see Materials and Methods for details). AP/MS was carried out using 0–3 hr embryos from mothers expressing *UAS-GFP-dTgs1* driven by *actin-Gal4* or *Smn-GFP* under the control of the tubulin promoter. (A) Efficiency of GFP-TRAP-mediated AP assayed using an anti-GFP antibody (T, total protein extract; I, input (10%); S supernatant; IP, immunoprecipitate). (B, C) dTgs1 (B) and Smn (C) interacting proteins; All protein IDs conforming to stringent filtering (see Materials and Methods) are shown in the Tables. Mean area corresponds to Top 3 protein quantification (T3PQ); the mean of the three highest abundance peptides identified for each protein. The complete lists of the Tgs1 and Smn interactors are shown in tables S1 and S2.

### Mutations in *dTgs1* cause neurological phenotypes

Previous studies have shown that loss-of-function mutations in the *Drosophila Smn* gene result in a variety of phenotypes, including alterations in the sensory-motor neuronal network, abnormal neuromuscular junctions, and defective locomotion [44,51-55]. In addition, we have recently shown that Smn depletion in neurons results in unexpanded wings and unretracted ptilinum [42]. The ptilinum is a head muscle required to break open the operculum of the puparium, and is normally retracted after fly eclosion. The post-eclosion events are regulated by the bursicon neuropeptide and a specific set of neurons [56,57], suggesting that the wing expansion/ptilinum phenotype could be a consequence of improper functioning of the underlying neural circuit. The finding that dTgs1 interacts with all *Drosophila* Smn components prompted us to ask whether its loss result in a phenotype comparable to that observed after Smn depletion. To analyze the role of *dTgs1* in *Drosophila* we took advantage of *moi*^*1*^, a point mutation within the *Drosophila moi-dTgs1* bicistronic locus that affects the function of both *moi* and *Tgs1* (Figure 1A, B). *moi*^*1*^ homozygotes are viable in the presence of a *moi*^*+*^ transgene, and both in the presence and in the absence of this transgene, they exhibit a substantially lower level of dTgs1 (∼ 50%) compared to *moi*^*1*^*/+* heterozygotes or *moi*^*1*^ homozygotes bearing the *dTgs1*^*FL*^ transgene (Figure 3A).

**Figure 3.**
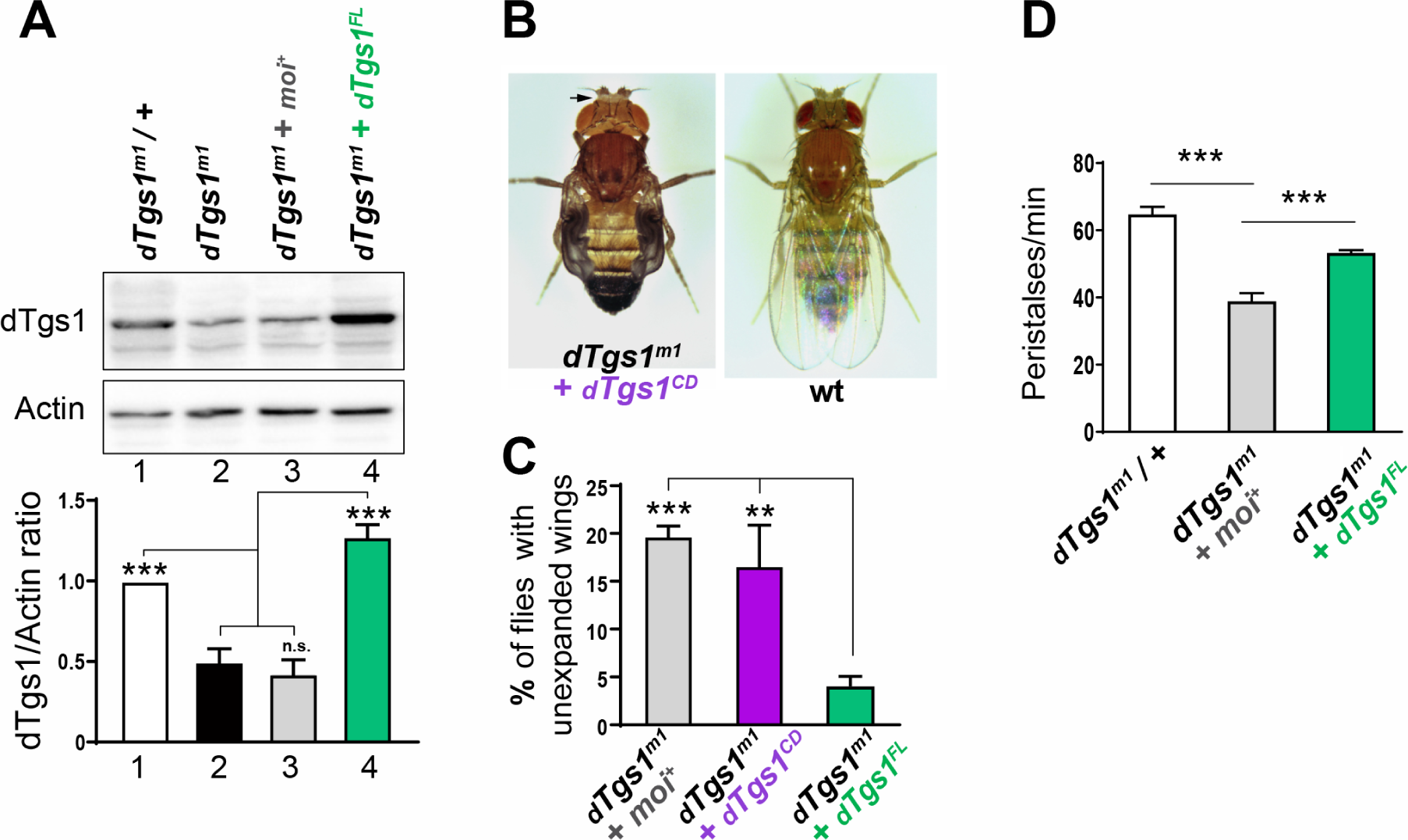
Flies homozygous for *dTgs1* hypomorphic mutations exhibit defective wing expansion and unretracted ptilinum. (A) Top panel: Representative Western blot showing the reduced abundance of the dTgs1 protein in larval brains of d*Tgs1*^*m1*^ homozygous flies in the presence or absence of the *moi*^*+*^ rescue construct (lanes 2 and 3), compared to d*Tgs1*^*m1*^ homozygotes bearing the *dTgs1*^*FL*^ rescue construct (lane 4) or d*Tgs1*^*m1*^*/+* heterozygous flies (control, lane 1). Bottom panel: quantification of the dTgs1 protein level relative to the actin loading control. Data are representative of 3 independent experiments, 10 larval brains per sample. ***, p< 0.001; one way ANOVA. (B) Defects in wing expansion and ptilinum retraction (arrow) observed in d*Tgs1*^*m1*^ homozygous flies bearing either the *moi*^*+*^ or the *dTgs1*^*CD*^ rescue construct (described in Figures 1A, B); wt, wild type. (C) Frequencies of wing expansion defects in d*Tgs1*^*m1*^ homozygous flies expressing the indicated rescue constructs: *moi*^*+*^ (n = 691), *dTgs1*^*CD*^ (n = 3802) and *dTgs1*^*FL*^ (n = 2744). **, p< 0.01; ***, p< 0.001; one-way ANOVA. (D) Frequencies of peristaltic contraction rates in larvae of the indicated genotypes (larvae bearing the moi^+^ or the *Tgs1*^*FL*^ construct are homozygous for d*Tgs1*^*m1*^). From left to right, 12, 10 and 12 larvae were analyzed. ***, p< 0.001; one-way ANOVA

Thus the *moi*^*1*^ mutation is also hypomorphic for *dTgs1* and will be henceforth designated as d*Tgs1*^*m1*^. We also exploited a rescue construct, *dTgs1*^*CD*^, encoding the entire *moi-dTgs1* sequence but carrying point mutations within the dTgs1 methyltransferase domain [9] (Figure 1A, B). *dTgs1*^*CD*^ rescued the lethality of *dTgs1*^*m1*^ homozygotes (Figure 1B), but did not complement the *dTgs1*^*R1*^ and *dTgs1*^*R2*^ null mutations, indicating that the dTgs1 protein encoded by this construct is not functional.

Adult flies homozygous for d*Tgs1*^*m1*^ and bearing either the *moi*^*+*^ or *dTgs1*^*CD*^ transgene showed a high proportion (20% and 16%, respectively) of individuals displaying defects in wing expansion and ptilinum retraction (Figure 3B, C). These defects were strongly reduced in flies homozygous for *dTgs1*^*m1*^ but bearing a *dTgs1*^*FL*^ transgene (Figure 3C). Thus, the wing expansion and the ptilinum phenotypes are caused by an impairment of the *dTgs1* function. We also investigated whether *dTgs1* mutant larvae exhibit locomotion defects, measured as frequency in contraction rates (peristalses). We found that *dTgs1*^*m1*^ homozygous larvae bearing a *moi*^*+*^ transgene exhibit a reduction in the frequency of peristalses compared to both *dTgs1*^m*1*^/+ heterozygotes and *dTgs1*^*m1*^ homozygotes bearing the *dTgs1*^*FL*^ rescue construct (Figure 3D).

### Tissue-specific silencing of dTgs1

To further explore the role of *dTgs1* we used flies bearing the transgenic construct *UAS-dTgs1 RNAi* (abbreviated as *dTgs1-RNAi*) already described by [37]. To check the efficiency of this RNAi construct we determined the dTgs1 protein level in larval brains carrying (i) *dTgs1-RNAi* and the *actin-Gal4* driver in a wild type background and (ii) *dTgs1-RNAi* and *actin-Gal4* in a *dTgs1*^*m1*^*/dTgs1*^*+*^ background; these brains showed 60% and 70% reduction in dTgs1 compared to control brains carrying *dTgs1-RNAi* and no driver, respectively (Figure 4A). The dTgs1 reduction was even stronger (∼ 80%) when RNAi against *dTgs1* was performed in flies heterozygous for the *dTgs1*^*R1*^ mutation (Figure 4B). To assess the role of dTgs1in different tissues, we crossed *dTgs1-RNAi*-bearing flies with flies carrying various Gal4 drivers with different tissue specificities (Figure 4C). The *dTgs1-RNAi* transgene in the presence of either the ubiquitously expressed *actin-Gal4* driver or the *how24B-Gal4* mesodermal driver (targeting muscles) induced lethality in third instar larval/early pupal stages (Figure 4C). The pan-neuronal drivers *elav155-Gal4* and *nsyb-Gal4* did not induce lethality or visible morphological phenotypes (Figure 4C), such as wing expansion defects, which were instead observed when a *UAS-Smn RNAi* construct is expressed by the same drivers [42]. The results of these RNAi experiments are in line with those previously reported by [37].

**Figure 4.**
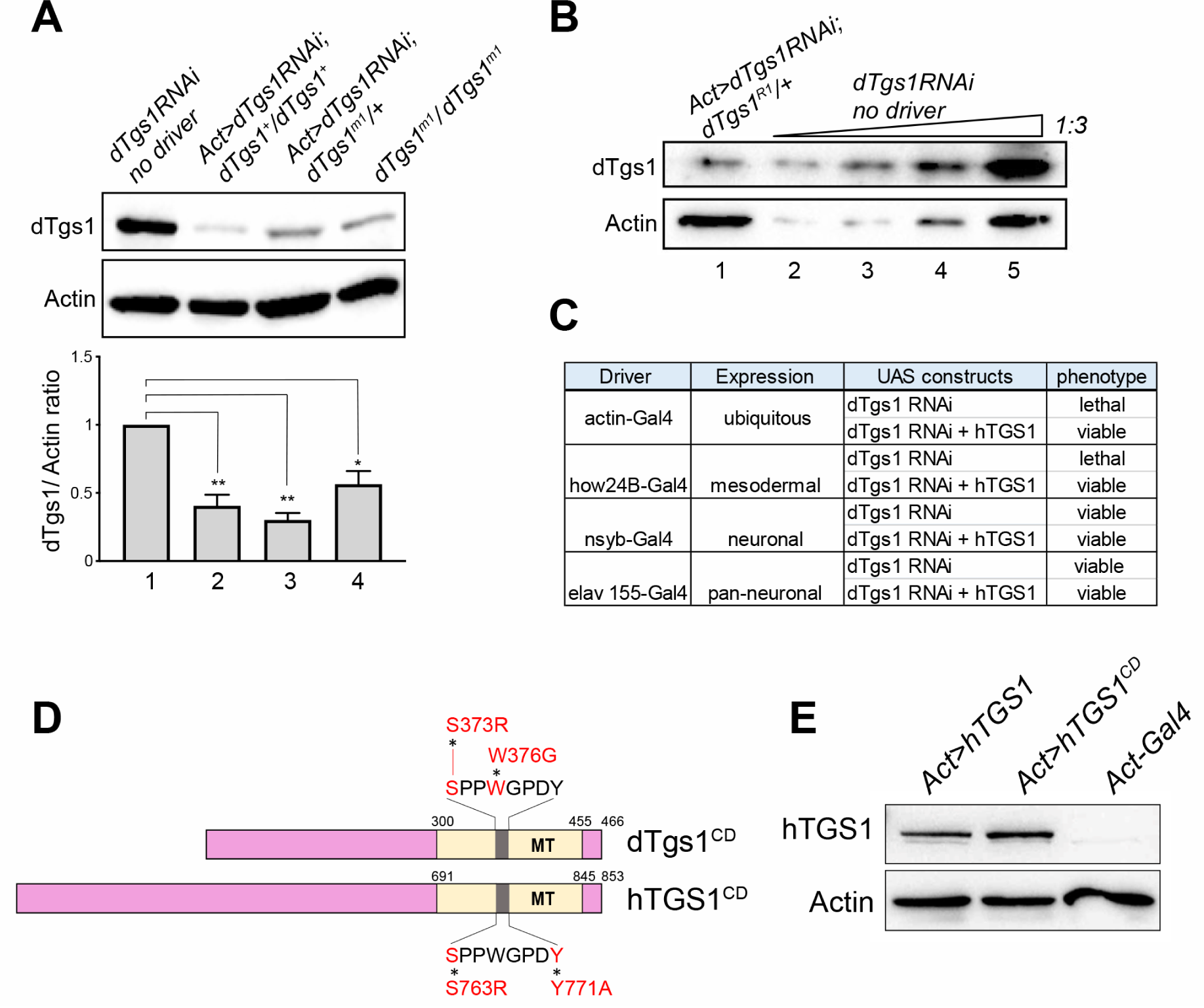
Human *TGS1* rescues the lethality induced by RNAi-mediated depletion of *dTgs1.* (A) Top panel: Representative western blot showing the dTgs1 levels in brains from larvae bearing the *UAS*-*dTgs1* RNAi construct (*dTgs1*RNAi) and no driver (lane1), *dTgs1*RNAi and the *actin-Gal4* driver (*Act*>*dTgs1RNAi*) in a *dTgs1*^*+*^ background (lane 2) or in a *dTgs1*^*m1*^*/+* background (lane 3), or homozygous for d*Tgs1*^*m1*^ (lane 4). Bottom panel: Quantification of the dTgs1 protein levels relative to the Actin loading control in brains from larvae of the indicated genotypes. Data are from three experiments, 10 larval brains per sample. *, p< 0.05; ** p< 0,01; one-way ANOVA. (B) Western blot showing the reduced level of dTgs1 in brains from larvae heterozygous for the *dTgs1*^*R1*^ mutation, bearing the *UAS-dTgs1RNAi* construct driven by Actin-*GAL4*. In lanes 2, 3 and 4, the amount of extract loaded is 1/27, 1/9 and 1/3 of that of lane 5, respectively. (C) Phenotypes induced by the expression of *UAS-dTgs1 RNAi* driven by the indicated *Gal4* drivers, in the presence or absence of a *UAS-hTGS1* construct. (D) Schematic representation of catalytic dead versions of *Drosophila Tgs1* (*dTgs1*^*CD*^) and human *TGS1* (*hTGS1*^*CD*^) proteins, carrying the indicated amino acid substitutions targeting the conserved amino acid stretch (grey), within the methyltransferase domain (MT, yellow). (E) Representative western blot showing that hTGS1 and hTGS1^CD^ are expressed at similar levels in brains from larvae bearing the *actin-Gal4* driver and either *UAS-hTGS1* (*Act*>*hTGS1*) or *UAS-hTGS1*^*CD*^ (*Act*>*hTGS1*^*CD*^). Protein extracts are from 10 larval brains; Actin was used as the loading control.

### A human *TGS1* transgene rescues the phenotypes elicited by mutations in Drosophila *Tgs1*

We next asked whether, and to what extent, a human *TGS1* gene (henceforth designated as *hTGS1*) has the ability to functionally substitute for its *Drosophila* homolog. We generated a fly line bearing a wild type human *TGS1* gene fused with the UAS promoter (*UAS-hTGS1)*, and a second line carrying a similar construct in which the human *TGS1* gene was mutated in the catalytic site (*UAS-hTGS1*^*CD*^; Figure 4 D, E). To generate hTGS1^CD^, we substituted the residues S763 and Y771 within the highly conserved Motif IV of the hTGS1 methyltransferase domain (Figure 4D) [58].

This amino acid stretch is important for substrate binding and identifies the TGS1 catalytic center. Specific mutations in the S and W residues of this motif suppress the TGS1 catalytic activity in yeast, humans and *Drosophila* [5,9,58]. Western blotting analysis showed that *UAS-hTGS1* and *UAS-hTGS1*^*CD*^ are expressed at similar levels in larval brains in the presence of an actin-Gal4 driver (Figure 4E). We then generated *dTgs1*^*R1*^ and *dTgs1*^*R2*^ homozygotes bearing the *actin-Gal4* driver and either *UAS-hTGS1* or *UAS-hTGS1*^*CD*^. The wild type *hTGS1* transgene, but not *hTGS1*^*CD*^, rescued the lethality of both *dTgs1* mutants (Figure 1B).

We next used the human *TGS1* transgene (which is not targeted by our *Drosophila* RNAi construct) to ask whether *dTgs1-RNAi* affects the expression of both *dTgs1* and *moi*. We constructed *actin-Gal4*>*dTgs1-RNAi* and *how24B-Gal4*>*dTgs1-RNAi* flies (see Figure 4C) bearing *UAS-hTGS1* and found that the expression of the human transgene fully rescues the lethality caused by dTgs1 depletion. We also analyzed brain cells from *actin-Gal4*>*dTgs1-RNAi* larvae heterozygous for the *dTgs1*^*R1*^ mutation. Although the brains of these larvae displayed an 80% reduction in the Tgs1 level (Figure 4B), they did not show telomeric fusions (200 cells examined). Collectively, these results confirm that *dTgs1* plays *a moi-*independent function. Most importantly, they demonstrate a remarkable evolutionary conservation of the *TGS1* function, and that the dTgs1 catalytic site is essential for fly viability.

### *The roles of dTgs1* and *Smn* in eye development

We next examined the role of *dTgs1* and *Smn* in the morphogenesis of the *Drosophila* compound eye, a remarkably patterned structure highly suitable for the analysis of cell viability and proliferation. To ascertain a possible role of *dTgs1* in the eye development we exploited the *dTgs1-RNAi* construct and the *eyeless-Gal4* driver (*ey-Gal4*), which efficiently drives expression of UAS-RNAi constructs in the eye imaginal disc [59]. *eyeless* is an essential *Drosophila Pax6* gene that specifically promotes growth of the eye imaginal disc and it is required for eye progenitor cell survival and proliferation [60,61].

When we expressed the *dTgs1-RNAi* construct under the control of the *ey-Gal4* driver, we found that 92% of the eyes (n = 1,280) from these RNAi flies display a reduction in size with defects ranging from total eye absence to severe or mild eye reduction. We subdivided the eyes into 5 classes, ranging from class 0 characterized by the complete absence of the eye to class 100 that comprises essentially normal eyes; the sizes of the eyes in classes 25, 50, and 75 were < 25%, < 50%, and < 75% the size of wild type eyes, respectively (see Figure 5A and Material and Methods for details on eye size estimation). To assess the specificity of the effects of *dTgs1* downregulation on eye development we generated flies bearing the *ey-Gal4* driver, *dTgs1-RNAi* and the RNAi-resistant *UAS-hTGS1* construct. We found that *hTGS1* expression rescues the eye phenotype (n= 1,698 eyes examined), abrogating the 0-50 size classes and inducing a substantial increase in the proportion of eyes with a regular size (from 8% to 53%) (Figure 5B). In contrast, the expression of *hTGS1*^*CD*^ did not mitigate the eye phenotype (n = 344 eyes) of *ey-Gal4*>*Tgs1 RNAi* flies (Figure 5B). These results strongly suggest that the defective eye development observed in *dTgs1* RNAi flies is specifically due to loss of the dTgs1 hypermethylase activity.

To asses the role of Smn in eye development we generated *ey-Gal4*> *Smn-*RNAi flies, using a previously characterized *UAS-Smn-RNAi* construct [42]. When this construct is expressed under the control of the *act-Gal4* driver, RNAi larvae die in the third instar stage, and the abundance of the Smn protein in larval brains is reduced to 30% of the wild type level [42]. We found that *ey-Gal4* >*Smn-RNAi*-induced *Smn* downregulation, in flies heterozygous for the *Smn* ^*X7*^ deficiency (that removes only the *Smn* gene), strongly affects eye development. Examination of 1,232 eyes from *Smn* RNAi flies did not reveal cases of complete eye absence, but 82% of the eyes were substantially reduced compared to wild type (12% class 25, 24% class 50 and 46% class 75) (Figure 5C). Interestingly, some of the eyes we included in the class 100 appeared slightly deformed and larger than the wild type eyes. In addition, 20% of the normal-sized eyes had an irregular surface and showed protruding areas. In both *ey-Gal4*>*Tgs1-RNAi* and *ey-Gal4*>*Smn-RNAi* flies displaying a reduction of the eye also the antennae were strongly affected. We observed a variety of defects ranging from an abnormal number of antennae (usually 1 or 3) to the apparent transformation of the eye tissue into head epidermis or antennal tissue. The degree of eye-to-antenna transformation ranged from some antennal tissue protruding from the eye to the complete transformation of an eye into an antenna (Figure 5D). Similar homeotic fate transformation phenotypes have been previously described in mutants causing cell proliferation defects in the eye imaginal disc and when eye specification genes such as *ey, twin of eyeless* (*toy*), *eyes absent* (*eya*), *sine oculis* (*so*) or *dachshund (dac)* are downregulated or misregulated [62-64].

**Figure 5.**
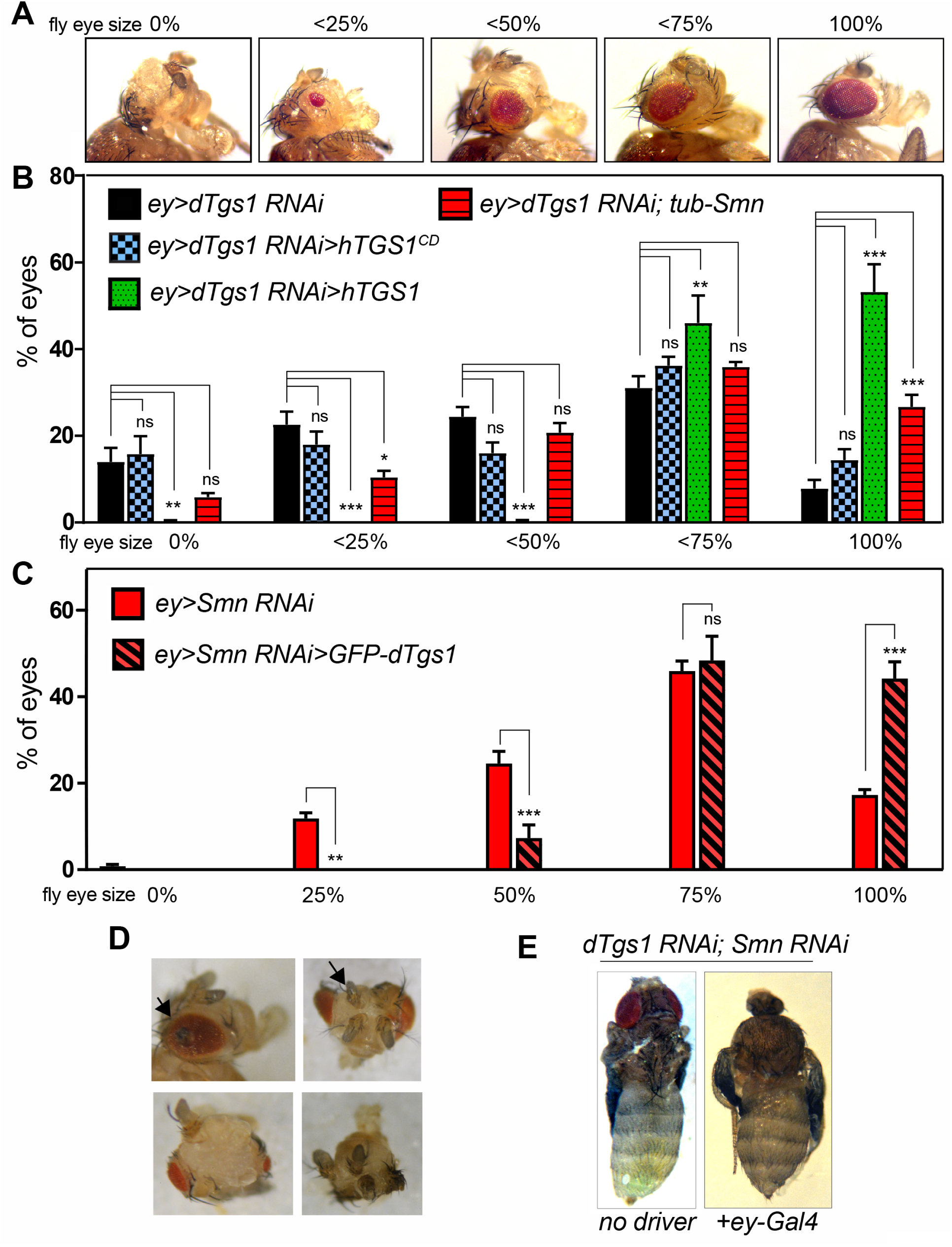
Genetic interactions between *dTgs1* and *Smn* in eye development. (A) Defects in eye development induced by *dTgs1* or *Smn* downregulation. The five panels show the range of defects induced by *eyeless-Gal4*-driven expression of *UAS-dTgs1RNAi* (*ey*>*Tgs1RNAi)* or *UAS-SmnRNAi* (*ey*>*SmnRNAi)*. Class 0%, eye absent; classes < 0.25%, < 0.50% and < 0.75%, eyes with sizes below 25%, 50% and 75% of the wild-type eye size (class100%). (B) Eye sizes in flies expressing *UAS-dTgs1 RNAi* and the indicated transgenes driven by *ey-Gal4* (ey>). The histogram bars represent the percentages of eyes falling into each size class shown in (A). The *ey*>*dTgs1RNAi*>*hTGS1* and *ey*>*dTgs1RNAi*> *hTGS1*^*CD*^ carry a *dTgs1RNAi* construct and either the *UAS-hTGS1* or the *UAS-hTGS1*^*CD*^ transgene, all driven by *ey-Gal4*; the *ey*>*dTgs1 RNAi*; *tub-*Smn flies express an *Smn-FLAG* transgene under control of the ubiquitous tubulin promoter. *, p< 0.05; **, p< 0.01; ***, p< 0.001; ns, not significant; two-way ANOVA. Note that the eye defects elicited by *dTgs1 RNAi* are partially rescued by *hTGS1* but not by *hTGS1*^*CD*^, and that the *dTgs1* mutant phenotype is ameliorated by Smn overexpression. (C) Eye sizes in flies carrying the *ey-Gal4* driver and either the *UAS-SmnRNAi* construct only (*ey*>*SmnRNAi*) or both *UAS-SmnRNAi* and *UAS-GFP-dTgs1.* The histogram bars represent the percentages of eyes falling into each size class shown in (A). ** p< 0,01, *** p< 0,001, ns, not significant; two-way ANOVA. Note that the eye defects elicited by *Smn RNAi* are partially rescued by overexpression of *dTgs1*. (D) Representative examples of partial eye-to-antenna transformation and antennal duplications observed in *ey*>*Tgs1RNAi* and in *ey*>*SmnRNAi* flies. (E) *ey-Gal4-*driven coexpression of the *dTgs1RNAi* and the *SmnRNAi* constructs induces synthetic lethality and defective head development. Left panel: control pharate adult fly manually extracted from the puparium with a fully developed head structure. Right panel: a *dTgs1 and Smn double RNAi* fly at a similar development stage lacking head structures.

To substantiate our data on the eye and antennal phenotypes we analyzed the eye-antennal imaginal discs from *ey-Gal4*>*Tgs1-RNAi* and *ey-Gal4*>*Smn-RNAi* third instar larvae. Fixed imaginal discs were stained with both anti-caspase and anti Elav antibodies and counterstained for DNA with DAPI. Elav staining identifies the photoreceptor neural cells placed posteriorly to the morphogenetic furrow [65] and Caspase-3 forms brightly fluorescent aggregates in correspondence to the apoptotic cells [66] (see Figure 6). We found that the eye imaginal discs of *ey-Gal4*>*Tgs1-RNAi* third instar larvae are significantly smaller (figure 6A, B) and exhibit a reduced proportion of Elav positive tissue compared to their wild type counterparts (figure 6A, C). In addition, these discs displayed an area enriched in apoptotic cells associated with bright caspase signals (figure 6A, D; see methods for quantitation of apoptotic signals); this area was located anteriorly to the morphogenetic furrow and involved retinal progenitor cells not stained by anti-Elav antibodies. We did not observe caspase signals in wild type discs. This is consistent with previous work [67] showing that apoptosis in the wild type eye disc does not occur in third instar larvae but only later in mid-pupal stages to eliminate the excess of pigment cells [68,69].

**Figure 6.**
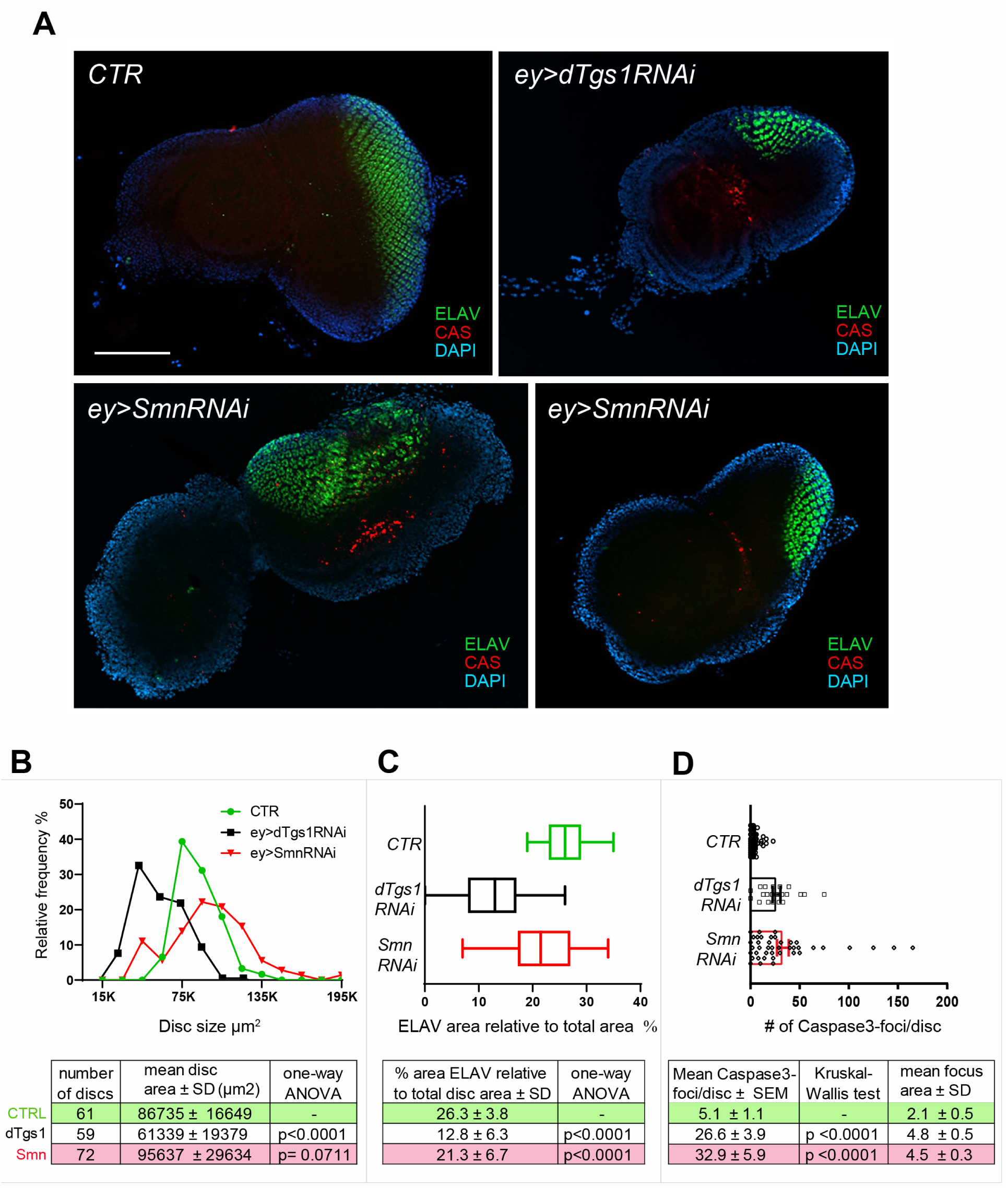
Downregulation of *dTgs1* or *Smn* in the eye imaginal discs induces apoptosis of retinal precursor cell and defective disc development. (A) Representative examples of eye-antennal imaginal discs from flies carrying *ey-Gal4* (*ey*) alone (CTR) or in combination with either the *dTgs1RNAi* or the *SmnRNAi* construct. Discs were stained for ELAV (which labels developing photoreceptor cells) and Caspase-3 (CAS, which marks cells undergoing apoptosis) and counterstained with DAPI. Scale bar: 100 μm. (B) Distributions of the size of the eye-antennal imaginal discs from flies with the genotypes indicated in (A). The number of discs analyzed, the mean disc area and descriptive statistics are reported in the graph below. (C) Box plots showing the quantification of the ELAV-labeled area in the discs described in (B), expressed as a percentage of the total disc area.The lines inside the boxes indicate the median; box boundaries represent the first and third quartiles; whiskers are 1.5 interquartile range. Descriptive statistics is reported below the graph. (D) Scatter dot-plots showing the quantification of the number of Caspase-3 positive foci per disc, in the discs described in (B). The mean area of the Caspase-3 foci and descriptive statistics are reported below the graph.

The imaginal discs from *ey-Gal4*>*Smn-RNAi* larvae showed defects that correlate with the eye phenotype observed in these animals. Indeed, the discs of these larvae showed a greater variability in size compared to the wild type discs (Figure 6B). *Smn* RNAi disc sizes had a normal distribution like that of wild type discs, but many of them had sizes that were either 2 standard deviations (SDs) below (11%) or above (19%) the mean size of control discs (Figure 6A, B). The *Smn* RNAi discs showed a modest but significant reduction in the proportion of Elav positive tissue compared to wild type discs (Figure 6A, C). They displayed areas with retinal progenitor cells undergoing apoptosis and did not show apoptotic signals associated with Elav-stained cells (Figure 6A, D). In summary, *Tgs1* or *Smn* knockdown in the eye imaginal disc results in apoptosis of retinal progenitor cells.

### Functional interactions between *dTgs1* and *Smn* in *Drosophila* eye development

To further explore the functional relationships between *dTgs1* and *Smn* we first performed a phenotypic analysis of flies with eye-targeted *dTgs1* silencing, simultaneously overexpressing *Smn*. We generated *ey-Gal4*>*dTgs1-RNAi* flies bearing a ubiquitously expressed *Smn-FLAG* construct under the control of the tubulin promoter (*tub-Smn*) described by [42]. A comparison of these flies (n = 1,248 eyes) with the *ey-GAL4*>*Tgs1-RNAi* flies (n = 1,280 eyes), or the *ey-Gal4*>*Tgs1-RNAi* flies expressing *hTGS1*^*CD*^ (n = 344 eyes), showed that the expression of Smn-FLAG substantially ameliorates their eye phenotype, reducing the percentage of very small eyes (classes 0 and 25) from 37% to 16% and increasing class 100 from 8% to 27% (Figure 5B).

We next analyzed the effect of *dTgs1* overexpression in Smn deficient-eyes. We constructed *ey-Gal4*>*Smn-RNAi* flies heterozygous for the *Smn*^*x7*^ deficiency and compared their eye phenotype (616 eyes) with that of flies of the same genetic constitution but bearing a *UAS-GFP-dTgs1* construct (and thus overexpressing dTgs1) (374 eyes). We found that overexpression of dTgs1 significantly counteracts the eye developmental defects caused by *Smn* silencing, reducing the 25 and 50 eye classes from 12% to 0% and from 25% to 7%, respectively, and increasing class 100 from 17% to 44% (figure 5C).

Finally, we analyzed flies bearing the *ey-Gal4* driver and both the *Tgs1-RNAi* and the *Smn-RNAi* constructs. 98% (n = 600) of these flies died at the pupal stage, and the rare surviving adults displayed severe eye defects. Interestingly, 90% of the *ey-GAL4*>*dTgs1-RNAi*>*Smn-RNAi* late lethal pupae manually extracted from the puparium showed a failure in development of the head structures (Figure 5E). Headless flies have been previously observed in mutants in genes that initiate eye specification such as the two fly Pax6 genes *ey* and *toy* [61,70]. Collectively, these results indicate that *dTgs1* and *Smn* cooperate in the pathways involved in eye and head development.

## Discussion

### The evolutionary conservation and the functional role of Tgs1

*Drosophila dTgs1* is part of a bicistronic locus that also includes *modigliani (moi)*, a gene required to prevent telomeric fusions (TFs). Previous complementation analyses with suitable transgenes suggested that the two genes play independent functions [8,9]. Here, we have provided a series of new data that support this conclusion. Specifically, we have used the CRISPR/Cas9 technology to introduce early stop codons in the *dTgs1* and *moi* coding sequences, and shown that these lethal mutations fully complement for viability and fertility. In addition, we have shown that the long isoform of the human *TGS1* transgene (*hTGS1*) fully rescues the lethality of flies homozygous for *dTgs1* null mutations. Notably, *hTGS1* also rescues the lethality caused by RNAi-mediated depletion of *dTgs1*, and rescued larvae do not exhibit TFs in brains. This result not only provides further support to the functional independence of *moi* and *dTgs1*, but clearly shows that RNAi against *dTgs1* does not disrupt the *moi* mRNA, validating the use of RNAi for specific impairment of the *dTgs1* function.

Previous work has shown that *TGS1* has high degree of functional conservation. The eukaryotic TGS1 proteins have different sizes and different degrees of global homology. However their C-terminal parts, that contain the methyltransferase catalytic domain, are structurally and functionally similar [58]. For example, loss of *S. cerevisiae TGS1* is complemented by a wild type human *TGS1* gene but not by a *TGS1* variant with mutations in the catalytic site [5]. Similarly, the growth inhibition phenotype caused by mutations in *S. cerevisiae TGS1* is rescued by a wild type *Arabidopsis TGS1* gene but not by a gene carrying mutations in the methyltransferase catalytic domain [6]. In line with these results, we have demonstrated that *dTgs1* and *hTGS1* carrying mutations in the catalytic domain (*dTgs1*^*CD*^ and *hTGS1*^*CD*^) are unable to rescue the phenotypic consequences of *dTgs1* deficiency. These results show that a human *TGS1* gene can substitute for its *Drosophila* ortholog. They also indicate that the lethality of *dTgs1* mutant and *dTgs1* RNAi flies, and the defective eye development elicited by targeted *dTgs1* depletion, are specifically due to loss of the dTgs1 hypermethylase activity.

### The biochemical relationships between Tgs1 and Smn

Previous studies on yeast and human cells have shown that TGS1 directly binds the SMN protein and associates with the SmB subunit of the Sm complex. It has been proposed that SMN recruits TGS1 favoring its engagement with the m7G-capped snRNA bound to the Sm ring, so as to allow hypermethylation of the cap [2,12]. The TMG cap and the Sm ring are thought to act as a bipartite nuclear-localization signal that mediates nuclear import of the snRNP particles [13,71-73]. Work in *Drosophila* has shown that that dTgs1 interacts with the Gemin3 subunit of the SMN complex in a two-hybrid assay [37]. Interestingly, the fly SMN complex contains only four bona fide Gemin proteins (Gem2, Gem3, Gem4/Glos and Gem5/Rig) [49,50,74]. The existence of potential homologs of Gem6, 7 and 8 was postulated by [50] but was not confirmed in a subsequent study [49]. Our AP/MS analyses of embryo extracts showed that Smn strongly co-precipitates with GFP-dTgs1. The other 11 most highly abundant dTgs1-interacting proteins included four Gemins (Gem2, Gem3, Gem4a and Gem4b). Conversely, the reciprocal AP-MS, using Smn-GFP as a bait, detected Gem3, Gem2, *Gem4a* and *Gem4b*, Lsm11, the cap binding protein Cbp80 and dTgs1 all within the top 11 interactors. Although Gem5 and other Sm complex subunits, such as SmD2 and SmD3, were not enriched in either AP-MS, they were present in the precipitates (Tables S1 and S2), and so specific biochemical associations between these proteins and Tgs1 or Smn cannot be ruled out; a physical interaction between TGS1 and SmB has previously been demonstrated in both human and yeast cells [12]. Moreover, consistent with our AP/MS results, Gem5 was the less abundant precipitate among the Gemins that co-purify with Smn-FLAG [49]. Collectively, our findings reveal a strong physical association between dTgs1 and the Smn complex subunits that has not been detected in previous studies on any organism. This suggests that dTgs1 and Smn may have intimate functional relationships.

### The functional relationships between Tgs1 and Smn

We have shown that homozygotes for *dTgs1* null mutation die in the second larval instar, and that that hypomorphic mutations in *dTgs1* are defective in in wing expansion and ptilinum retraction, a phenotype that likely reflects a perturbation in the neural circuit that coordinates the post-eclosion performance [56,57]. We also found that third instar larvae homozygous for *dTgs1*^*m1*^ hypomorphic mutations exhibit a reduction in the frequency of peristalses compared to controls. These results are consistent with previous studies showing that *dTgs1* mutant or RNAi flies exhibit abnormal larval and adult locomotor behaviors [8,37].

Locomotion phenotypes similar to those observed in *dTgs1* mutants have been previously seen in flies carrying mutations in *Smn* or expressing RNAi constructs against *Smn*. The unexpanded wing phenotype has been also observed flies where *Smn* was specifically silenced in neurons using RNAi [42]. *Smn* mutant larvae are defective in the sensory-motor neuronal network and exhibit reduced muscle growth and defective locomotion [44,54,55,75]. These phenotypes have been attributed to reduced biogenesis of snRNAs leading to defective splicing of a subset U12 intron-containing RNAs, altering the expression of genes required for motor circuit function such as *Stasimon* [20]. Other studies suggested that splicing defects in other RNA types such as those encoded by genes involved in stress response could contribute to the fly neurological phenotype [21,76]. Studies in *Drosophila* have also shown that the Smn function is not restricted to motor neurons. For example, Smn is required for stem cell division and differentiation [31], and for maintenance of proper organization of nuclear compartments in both nurse cells and oocytes [77]. Whether these functions of Smn depend on its role in snRNP biogenesis and splicing is currently unclear.

Here, we have documented another Smn function not involved in motor neuron maintenance. We have shown that both *Smn* and *dTgs1* are required for *Drosophila* eye development. *eyeless-Gal4*-driven RNAi against each of these genes resulted in high frequencies of flies with reduced eye sizes. RNAi against *dTgs1* produced small eye discs containing a reduced proportion of Elav-positive differentiated cells compared to wild type. The eye discs of *Smn* RNAi flies were greatly variable in size; some of them were bigger and other smaller than their wild type counterparts. They also showed a modest but significant reduction of the proportion of Elav-positive cells. Both *dTgs1* and *Smn* RNAi discs displayed frequent apoptotic cells in the anterior disc area that contain undifferentiated retinal progenitor cells but not in the posterior area containing Elav-stained cells. These results suggest that both d*Tgs1* and Smn are required to prevent cell damage leading to apoptosis of retinal progenitor cells. When either gene is downregulated, precursor cells die leading to a reduction in the differentiated retinal cells and to an overall reduction of the eye size. The few large discs and eyes observed after *Smn* RNAi might be the consequence of compensatory proliferation or apoptosis-induced proliferation of imaginal cells, two phenomena that compensate for disc cell loss or damage [78,79]and references therein).

We have also demonstrated that *Smn* and *dTgs1* interact genetically in the control of eye disc development. Overexpression of *Smn* substantially ameliorates the eye phenotype caused by dTgs1 depletion, while *dTgs1* overexpression partially rescues the eye defects elicited by *Smn* downregulation. In addition, double RNAi against both *dTgs1* and *Smn* greatly exacerbates the phenotypes observed in single RNAi flies, leading to severe eye defects and frequent headless individuals unable to emerge from the pupal case. These results suggest that *dTgs1* and *Smn* cooperate in a pathway that controls survival of retinal precursor cells. We do not know the precise roles of these genes in this pathway but hypothesize that eye mutant phenotypes are the consequence of defective snRNP biogenesis and pre-mRNA splicing. There is ample evidence that Smn controls these processes [47,80], and we have recently found that *Drosophila* cells carrying mutations in *dTgs1* accumulate snRNAs with defective 3’ terminations (our unpublished results). Since the protein products of *dTgs1* and *Smn* strongly interact, we can postulate that overexpression of one of the genes can partially compensate for downregulation of the other, and that inhibition of both genes can worsen the situation. However, assuming that both genes control pre-mRNA splicing in the eye imaginal disc, their specific pre-mRNA targets remain to be determined.

### The *Drosophila* eye as model system for identifying modifiers of the *Tgs1* and *Smn* loss of function phenotypes

The *Drosophila* model has been extremely useful to identify modifiers of the *Smn* loss of function phenotype. As mentioned earlier, molecular analysis of *Drosophila Smn* mutants led to the identification of *Stasimon*, an *Smn* target gene whose expression in neurons restores the neurological defects elicited by Smn mutations [20]. Stasimon orthologs share a conserved function and ameliorate the *Smn* loss of function phenotype also in zebrafish and mice [20,25]. Other modifiers of the *Smn*-dependent phenotype have been isolated for their interaction with hypomorphic *Smn* mutations. For example, combinations of a specific *Smn* hypomorphic allele a with a large number of insertional mutations affecting approximately 50% of the fly genes (the Exelixis collection) led to the identification of 17 enhancers and 10 suppressors of the *Smn* mutant phenotype, most of which affected the neuromuscular junction (NMJ) [44]. The *C. elegans* orthologs of 12 of these genes were able to modify the *Smn* loss of function defects in the worm [81]. The same study showed that the invertebrate orthologs of *Plastin 3* (*PLS3*), a human SMA modifier, also modify of the *Smn* loss of function phenotype in both *C. elegans* and *Drosophila* models. More recently, we analyzed the roles of the *Drosophila* and *C. elegans* orthologs of the human *WDR79 /TCAB1* gene, which encodes a protein that interacts with SMN and controls the biogenesis of several RNA species. Downregulation of these genes in flies and worms resulted in locomotion defects similar to those elicited by Smn depletion. In addition, we found that WDR79 ameliorates the locomotion defects caused by Smn depletion and vice versa [42]. Collectively these results indicate that the *Smn* interacting genes identified in *Drosophila* are conserved across species, and reinforce the idea that *Drosophila* is a well-suited model organism for detecting *Smn* loss of function modifiers, providing potentially helpful information for SMA therapy.

We have recently shown that *TGS1* mutations cause an increase in the hTR level and telomerase activity in human cells, leading to telomere elongation [4]. There is currently a great interest in conceiving new therapies for diseases caused by telomerase insufficiency such as dyskeratosis congenita and disorders associated with human aging. The *dTgs1* loss of function eye model could provide an important contribution to the research in this area, as it allows an easy detection of chemical and genetic modifiers of the eye phenotype. Identification of such modifiers might help devising new approaches to the control of human telomere length.

## Acknowledgements

We thank Spyros Artavanis-Tsakonas for fly strains, and Ammarah Tariq for assistance with the GFP-dTgs1 affinity purification. This work was supported by grants from Telethon (GPP13147) and Fondazione Cenci Bolognetti to G.D.R., and Associazione Italiana Ricerca sul Cancro (AIRC, IG 20528) to M.G. G.N. was supported by fellowships from Regione Lazio (Torno Subito 2018) and from Sapienza, University of Rome.

## References

1. Girard C, Verheggen C, Neel H, Cammas A, Vagner S, et al. (2008) Characterization of a short isoform of human Tgs1 hypermethylase associating with small nucleolar ribonucleoprotein core proteins and produced by limited proteolytic processing. J Biol Chem 283: 2060–2069.

2. Mouaikel J, Verheggen C, Bertrand E, Tazi J, Bordonne R (2002) Hypermethylation of the cap structure of both yeast snRNAs and snoRNAs requires a conserved methyltransferase that is localized to the nucleolus. Mol Cell 9: 891–901.

3. Wurth L, Gribling-Burrer AS, Verheggen C, Leichter M, Takeuchi A, et al. (2014) Hypermethylated-capped selenoprotein mRNAs in mammals. Nucleic Acids Res 42: 8663–8677.

4. Chen L, Roake CM, Galati A, Bavasso F, Micheli E, et al. (2020) Loss of Human TGS1 Hypermethylase Promotes Increased Telomerase RNA and Telomere Elongation. Cell Rep 30: 1358–1372 e1355.

5. Hausmann S, Zheng S, Costanzo M, Brost RL, Garcin D, et al. (2008) Genetic and biochemical analysis of yeast and human cap trimethylguanosine synthase: functional overlap of 2,2,7-trimethylguanosine caps, small nuclear ribonucleoprotein components, pre-mRNA splicing factors, and RNA decay pathways. J Biol Chem 283: 31706–31718.

6. Gao J, Wallis JG, Jewell JB, Browse J (2017) Trimethylguanosine Synthase1 (TGS1) Is Essential for Chilling Tolerance. Plant Physiol 174: 1713–1727.

7. Komonyi O, Papai G, Enunlu I, Muratoglu S, Pankotai T, et al. (2005) DTL, the Drosophila homolog of PIMT/Tgs1 nuclear receptor coactivator-interacting protein/RNA methyltransferase, has an essential role in development. J Biol Chem 280: 12397–12404.

8. Komonyi O, Schauer T, Papai G, Deak P, Boros IM (2009) A product of the bicistronic Drosophila melanogaster gene CG31241, which also encodes a trimethylguanosine synthase, plays a role in telomere protection. J Cell Sci 122: 769–774.

9. Raffa GD, Siriaco G, Cugusi S, Ciapponi L, Cenci G, et al. (2009) The Drosophila modigliani (moi) gene encodes a HOAP-interacting protein required for telomere protection. Proceedings of the National Academy of Sciences of the United States of America 106: 2271–2276.

10. Jia Y, Viswakarma N, Crawford SE, Sarkar J, Sambasiva Rao M, et al. (2012) Early embryonic lethality of mice with disrupted transcription cofactor PIMT/NCOA6IP/Tgs1 gene. Mech Dev 129: 193–207.

11. Ohno M, Segref A, Bachi A, Wilm M, Mattaj IW (2000) PHAX, a mediator of U snRNA nuclear export whose activity is regulated by phosphorylation. Cell 101: 187–198.

12. Mouaikel J, Narayanan U, Verheggen C, Matera AG, Bertrand E, et al. (2003) Interaction between the small-nuclear-RNA cap hypermethylase and the spinal muscular atrophy protein, survival of motor neuron. EMBO Rep 4: 616–622.

13. Gruss OJ, Meduri R, Schilling M, Fischer U (2017) UsnRNP biogenesis: mechanisms and regulation. Chromosoma 126: 577–593.

14. Carissimi C, Saieva L, Gabanella F, Pellizzoni L (2006) Gemin8 is required for the architecture and function of the survival motor neuron complex. J Biol Chem 281: 37009–37016.

15. Franke J, Gehlen J, Ehrenhofer-Murray AE (2008) Hypermethylation of yeast telomerase RNA by the snRNA and snoRNA methyltransferase Tgs1. J Cell Sci 121: 3553–3560.

16. Tang W, Kannan R, Blanchette M, Baumann P (2012) Telomerase RNA biogenesis involves sequential binding by Sm and Lsm complexes. Nature 484: 260–264.

17. Beattie CE, Kolb SJ (2018) Spinal muscular atrophy: Selective motor neuron loss and global defect in the assembly of ribonucleoproteins. Brain Res 1693: 92–97.

18. Ruggiu M, McGovern VL, Lotti F, Saieva L, Li DK, et al. (2012) A role for SMN exon 7 splicing in the selective vulnerability of motor neurons in spinal muscular atrophy. Mol Cell Biol 32: 126–138.

19. Zhang Z, Lotti F, Dittmar K, Younis I, Wan L, et al. (2008) SMN deficiency causes tissue-specific perturbations in the repertoire of snRNAs and widespread defects in splicing. Cell 133: 585–600.

20. Lotti F, Imlach WL, Saieva L, Beck ES, Hao le T, et al. (2012) An SMN-dependent U12 splicing event essential for motor circuit function. Cell 151: 440–454.

21. Garcia EL, Wen Y, Praveen K, Matera AG (2016) Transcriptomic comparison of Drosophila snRNP biogenesis mutants reveals mutant-specific changes in pre-mRNA processing: implications for spinal muscular atrophy. RNA 22: 1215–1227.

22. Rizzo F, Nizzardo M, Vashisht S, Molteni E, Melzi V, et al. (2019) Key role of SMN/SYNCRIP and RNA-Motif 7 in spinal muscular atrophy: RNA-Seq and motif analysis of human motor neurons. Brain 142: 276–294.

23. Van Alstyne M, Simon CM, Sardi SP, Shihabuddin LS, Mentis GZ, et al. (2018) Dysregulation of Mdm2 and Mdm4 alternative splicing underlies motor neuron death in spinal muscular atrophy. Genes Dev 32: 1045–1059.

24. Simon CM, Dai Y, Van Alstyne M, Koutsioumpa C, Pagiazitis JG, et al. (2017) Converging Mechanisms of p53 Activation Drive Motor Neuron Degeneration in Spinal Muscular Atrophy. Cell Rep 21: 3767–3780.

25. Simon CM, Van Alstyne M, Lotti F, Bianchetti E, Tisdale S, et al. (2019) Stasimon Contributes to the Loss of Sensory Synapses and Motor Neuron Death in a Mouse Model of Spinal Muscular Atrophy. Cell Rep 29: 3885–3901 e3885.

26. McWhorter ML, Monani UR, Burghes AH, Beattie CE (2003) Knockdown of the survival motor neuron (Smn) protein in zebrafish causes defects in motor axon outgrowth and pathfinding. J Cell Biol 162: 919–931.

27. Fallini C, Donlin-Asp PG, Rouanet JP, Bassell GJ, Rossoll W (2016) Deficiency of the Survival of Motor Neuron Protein Impairs mRNA Localization and Local Translation in the Growth Cone of Motor Neurons. J Neurosci 36: 3811–3820.

28. Donlin-Asp PG, Bassell GJ, Rossoll W (2016) A role for the survival of motor neuron protein in mRNP assembly and transport. Curr Opin Neurobiol 39: 53–61.

29. Donlin-Asp PG, Rossoll W, Bassell GJ (2017) Spatially and temporally regulating translation via mRNA-binding proteins in cellular and neuronal function. FEBS Lett 591: 1508–1525.

30. Kariya S, Obis T, Garone C, Akay T, Sera F, et al. (2014) Requirement of enhanced Survival Motoneuron protein imposed during neuromuscular junction maturation. J Clin Invest 124: 785–800.

31. Grice SJ, Liu JL (2011) Survival motor neuron protein regulates stem cell division, proliferation, and differentiation in Drosophila. PLoS Genet 7: e1002030.

32. Chang WF, Xu J, Chang CC, Yang SH, Li HY, et al. (2015) SMN is required for the maintenance of embryonic stem cells and neuronal differentiation in mice. Brain Struct Funct 220: 1539–1553.

33. Walker MP, Rajendra TK, Saieva L, Fuentes JL, Pellizzoni L, et al. (2008) SMN complex localizes to the sarcomeric Z-disc and is a proteolytic target of calpain. Hum Mol Genet 17: 3399–3410.

34. Anderton RS, Meloni BP, Mastaglia FL, Boulos S (2013) Spinal muscular atrophy and the antiapoptotic role of survival of motor neuron (SMN) protein. Mol Neurobiol 47: 821–832.

35. Zhao DY, Gish G, Braunschweig U, Li Y, Ni Z, et al. (2016) SMN and symmetric arginine dimethylation of RNA polymerase II C-terminal domain control termination. Nature 529: 48–53.

36. Kannan A, Bhatia K, Branzei D, Gangwani L (2018) Combined deficiency of Senataxin and DNA-PKcs causes DNA damage accumulation and neurodegeneration in spinal muscular atrophy. Nucleic Acids Res 46: 8326–8346.

37. Borg RM, Fenech Salerno B, Vassallo N, Bordonne R, Cauchi RJ (2016) Disruption of snRNP biogenesis factors Tgs1 and pICln induces phenotypes that mirror aspects of SMN-Gemins complex perturbation in Drosophila, providing new insights into spinal muscular atrophy. Neurobiol Dis 94: 245–258.

38. Port F, Chen HM, Lee T, Bullock SL (2014) Optimized CRISPR/Cas tools for efficient germline and somatic genome engineering in Drosophila. Proc Natl Acad Sci U S A 111: E2967–2976.

39. Gratz SJ, Cummings AM, Nguyen JN, Hamm DC, Donohue LK, et al. (2013) Genome engineering of Drosophila with the CRISPR RNA-guided Cas9 nuclease. Genetics 194: 1029–1035.

40. Bischof J, Maeda RK, Hediger M, Karch F, Basler K (2007) An optimized transgenesis system for Drosophila using germ-line-specific phiC31 integrases. Proc Natl Acad Sci U S A 104: 3312–3317.

41. Bruckner K, Kockel L, Duchek P, Luque CM, Rorth P, et al. (2004) The PDGF/VEGF receptor controls blood cell survival in Drosophila. Dev Cell 7: 73–84.

42. Di Giorgio ML, Esposito A, Maccallini P, Micheli E, Bavasso F, et al. (2017) WDR79/TCAB1 plays a conserved role in the control of locomotion and ameliorates phenotypic defects in SMA models. Neurobiology of Disease 105: 42–50.

43. Dietzl G, Chen D, Schnorrer F, Su KC, Barinova Y, et al. (2007) A genome-wide transgenic RNAi library for conditional gene inactivation in Drosophila. Nature 448: 151–156.

44. Chang HC, Dimlich DN, Yokokura T, Mukherjee A, Kankel MW, et al. (2008) Modeling spinal muscular atrophy in Drosophila. PLoS One 3: e3209.

45. Lattao R, Bonaccorsi S, Guan X, Wasserman SA, Gatti M (2011) Tubby-tagged balancers for the Drosophila X and second chromosomes. Fly (Austin) 5: 369–370.

46. Palumbo V, Pellacani C, Heesom KJ, Rogala KB, Deane CM, et al. (2015) Misato Controls Mitotic Microtubule Generation by Stabilizing the TCP-1 Tubulin Chaperone Complex [corrected]. Curr Biol 25: 1777–1783.

47. Raimer AC, Gray KM, Matera AG (2017) SMN - A chaperone for nuclear RNP social occasions? RNA Biol 14: 701–711.

48. Lemm I, Girard C, Kuhn AN, Watkins NJ, Schneider M, et al. (2006) Ongoing U snRNP biogenesis is required for the integrity of Cajal bodies. Mol Biol Cell 17: 3221–3231.

49. Matera AG, Raimer AC, Schmidt CA, Kelly JA, Droby GN, et al. (2019) Composition of the Survival Motor Neuron (SMN) Complex in Drosophila melanogaster. G3 (Bethesda) 9: 491–503.

50. Lanfranco M, Cacciottolo R, Borg RM, Vassallo N, Juge F, et al. (2017) Novel interactors of the Drosophila Survival Motor Neuron (SMN) Complex suggest its full conservation. FEBS Lett 591: 3600–3614.

51. Sen A, Yokokura T, Kankel MW, Dimlich DN, Manent J, et al. (2011) Modeling spinal muscular atrophy in Drosophila links Smn to FGF signaling. J Cell Biol 192: 481–495.

52. Praveen K, Wen Y, Gray KM, Noto JJ, Patlolla AR, et al. (2014) SMA-Causing Missense Mutations in Survival motor neuron (Smn) Display a Wide Range of Phenotypes When Modeled in Drosophila. PLoS Genet 10: e1004489.

53. Praveen K, Wen Y, Matera AG (2012) A Drosophila model of spinal muscular atrophy uncouples snRNP biogenesis functions of survival motor neuron from locomotion and viability defects. Cell Rep 1: 624–631.

54. Imlach WL, Beck ES, Choi BJ, Lotti F, Pellizzoni L, et al. (2012) SMN is required for sensory-motor circuit function in Drosophila. Cell 151: 427–439.

55. Spring AM, Raimer AC, Hamilton CD, Schillinger MJ, Matera AG (2019) Comprehensive Modeling of Spinal Muscular Atrophy in Drosophila melanogaster. Front Mol Neurosci 12: 113.

56. Luan H, Lemon WC, Peabody NC, Pohl JB, Zelensky PK, et al. (2006) Functional dissection of a neuronal network required for cuticle tanning and wing expansion in Drosophila. J Neurosci 26: 573–584.

57. Loveall BJ, Deitcher DL (2010) The essential role of bursicon during Drosophila development. BMC Dev Biol 10: 92.

58. Mouaikel J, Bujnicki JM, Tazi J, Bordonne R (2003) Sequence-structure-function relationships of Tgs1, the yeast snRNA/snoRNA cap hypermethylase. Nucleic Acids Res 31: 4899–4909.

59. Fan Y, Wang S, Hernandez J, Yenigun VB, Hertlein G, et al. (2014) Genetic models of apoptosis-induced proliferation decipher activation of JNK and identify a requirement of EGFR signaling for tissue regenerative responses in Drosophila. PLoS Genet 10: e1004131.

60. Callaerts P, Leng S, Clements J, Benassayag C, Cribbs D, et al. (2001) Drosophila Pax-6/eyeless is essential for normal adult brain structure and function. J Neurobiol 46: 73–88.

61. Zhu J, Palliyil S, Ran C, Kumar JP (2017) Drosophila Pax6 promotes development of the entire eye-antennal disc, thereby ensuring proper adult head formation. Proc Natl Acad Sci U S A 114: 5846–5853.

62. Kumar JP, Moses K (2001) EGF receptor and Notch signaling act upstream of Eyeless/Pax6 to control eye specification. Cell 104: 687–697.

63. Duong HA, Wang CW, Sun YH, Courey AJ (2008) Transformation of eye to antenna by misexpression of a single gene. Mech Dev 125: 130–141.

64. Kumar JP (2018) The fly eye: Through the looking glass. Dev Dyn 247: 111–123.

65. Koushika SP, Lisbin MJ, White K (1996) ELAV, a Drosophila neuron-specific protein, mediates the generation of an alternatively spliced neural protein isoform. Curr Biol 6: 1634–1641.

66. Fan Y, Bergmann A (2010) The cleaved-Caspase-3 antibody is a marker of Caspase-9-like DRONC activity in Drosophila. Cell Death Differ 17: 534–539.

67. Fan Y, Bergmann A (2008) Distinct mechanisms of apoptosis-induced compensatory proliferation in proliferating and differentiating tissues in the Drosophila eye. Dev Cell 14: 399–410.

68. Miller DT, Cagan RL (1998) Local induction of patterning and programmed cell death in the developing Drosophila retina. Development 125: 2327–2335.

69. Wolff T, Ready DF (1991) Cell death in normal and rough eye mutants of Drosophila. Development 113: 825–839.

70. Jiao R, Daube M, Duan H, Zou Y, Frei E, et al. (2001) Headless flies generated by developmental pathway interference. Development 128: 3307–3319.

71. Hamm J, Darzynkiewicz E, Tahara SM, Mattaj IW (1990) The trimethylguanosine cap structure of U1 snRNA is a component of a bipartite nuclear targeting signal. Cell 62: 569–577.

72. Narayanan U, Achsel T, Luhrmann R, Matera AG (2004) Coupled in vitro import of U snRNPs and SMN, the spinal muscular atrophy protein. Mol Cell 16: 223–234.

73. Natalizio AH, Matera AG (2013) Identification and characterization of Drosophila Snurportin reveals a role for the import receptor Moleskin/importin-7 in snRNP biogenesis. Mol Biol Cell 24: 2932–2942.

74. Cauchi RJ, Sanchez-Pulido L, Liu JL (2010) Drosophila SMN complex proteins Gemin2, Gemin3, and Gemin5 are components of U bodies. Exp Cell Res 316: 2354–2364.

75. Chan YB, Miguel-Aliaga I, Franks C, Thomas N, Trulzsch B, et al. (2003) Neuromuscular defects in a Drosophila survival motor neuron gene mutant. Hum Mol Genet 12: 1367–1376.

76. Garcia EL, Lu Z, Meers MP, Praveen K, Matera AG (2013) Developmental arrest of Drosophila survival motor neuron (Smn) mutants accounts for differences in expression of minor intron-containing genes. RNA 19: 1510–1516.

77. Lee L, Davies SE, Liu JL (2009) The spinal muscular atrophy protein SMN affects Drosophila germline nuclear organization through the U body-P body pathway. Dev Biol 332: 142–155.

78. Mollereau B, Perez-Garijo A, Bergmann A, Miura M, Gerlitz O, et al. (2013) Compensatory proliferation and apoptosis-induced proliferation: a need for clarification. Cell Death Differ 20: 181.

79. Su TT (2015) Non-autonomous consequences of cell death and other perks of being metazoan. AIMS Genet 2: 54–69.

80. Li DK, Tisdale S, Lotti F, Pellizzoni L (2014) SMN control of RNP assembly: from post-transcriptional gene regulation to motor neuron disease. Semin Cell Dev Biol 32: 22–29.

81. Dimitriadi M, Sleigh JN, Walker A, Chang HC, Sen A, et al. (2010) Conserved genes act as modifiers of invertebrate SMN loss of function defects. PLoS Genet 6: e1001172.

